# Decoding the mouse spinal cord locomotor neural network using tissue clearing, tissue expansion and tiling light sheet microscopy techniques

**DOI:** 10.1101/2022.07.04.498760

**Authors:** Ruili Feng, Jiongfang Xie, Jing Lu, Huijie Hu, Yanlu Chen, Dongyue Wang, Liang Gao

## Abstract

Decoding a biological neural network requires the structural information regarding the spatial organization, dendritic morphology, axonal projection and synaptic connection of the neurons in the network. Imaging physically sectioned nervous tissues using electron microscopy (EM) has been the only method to acquire such information. However, EM is inefficient for imaging and reconstructing large neural networks due to the low throughput and inability to target neural circuits of interest by labeling specific neuron populations genetically. Here, we present a method to image large nervous tissues from the cellular to synaptic level with high throughput using tiling light sheet microscopy combined with tissue clearing and tissue expansion techniques. We describe the method, demonstrate its capability and explore its utility for decoding large biological neural networks by studying the spinal cord locomotor neural network in genetically labeled fluorescent mice. We show our method could advance the decoding of large neural networks significantly.

## Introduction

Neurons acquire intellectual abilities by forming neural networks. In which neurons organize into multiple layers and communicate to each other via inter and intra layer connections (1–3). Thus, in addition to the neuron properties, the organization and connection of neurons determine the intellectual function of neural networks (4–7). As mammals are extremely capable of handling complicated tasks, it has been a long-term interest to understand the logical architecture and the underlying mathematical algorithm of the neural networks embodied in their central nervous system (CNS) (8–10). As with deriving the electric circuit schematic diagram by reverse engineering printed circuit boards. Outlining the connection diagram of neural networks is a necessary step to understand their intellectual function (11, 12). The structural information regarding the spatial organization, dendritic morphology, axonal projection and synaptic connection of the neurons in a neural network, which determines the neural network logical architecture and how information is exchanged and processed in the network, is required to accomplish the goal.

Imaging physically sectioned nervous tissues using electron microscopy (EM) has been an essential method to acquire the structural information of neural networks from the cellular to synaptic level for decades (13–22). However, EM suffers from the low throughput for imaging large tissues. Worse yet, the image contrast of EM is unrelated to specific neuron classes or neural circuits, which makes the interpretation of EM images for neural network diagrams a tougher challenge than the imaging itself. Therefore, visualizing and decoding biological neural networks using EM has been more successful on small model organisms like *C. elegans* and *Drosophila* (13,18,19) than on large organisms like rodents and primates. Furthermore, understanding a biological neural network requires observing the same network under different conditions, such as in individuals with different genetics, at different developmental stages and have different training experiences. Obviously, tissue imaging using EM will not satisfy the need in the near future.

Imaging nervous tissue slices using fluorescence microscopy (FM) is more practical in visualizing large neural networks due to its lower adoption threshold and higher throughput (23–32). More importantly, the utility of various genetic fluorescent labeling tools enables much more convenient observation of the neuron populations and neural circuits of interest (33–38). Unfortunately, besides the throughput of imaging tissue slices with FM still being low, caused by the cumbersome sample preparation, tissue sectioning and image registration of tissue slices, the spatial resolution is limited to the micron level which is insufficient to resolve complicated neuronal structures and synaptic connections between neurons.

The combination of tissue clearing (39–50), tissue expansion (51–58) and light sheet microscopy (LSM) (59–65) techniques offers an opportunity for a solution. The combination not only makes the 3D fluorescence imaging of nervous tissues and the following image reconstruction much more practical, but also provides sufficient yet flexible spatial resolving ability to observe biological neural networks from the cellular to synaptic level. Recently, the combination of tissue expansion techniques and lattice light sheet microscopy has been used to study neural circuits in *Drosophila*, proving the invaluable potential of the combination for understanding biological neural networks (56, 57). However, despite the successful implementations of the combination on small model organisms, its utility for studying large neural networks such as the mouse CNS neural networks remains challenging.

To respond to the challenge, we developed a tiling light sheet (TLS) microscope for imaging cleared and expanded tissues in our previous research (65–67). The microscope is compatible with most tissue clearing and tissue expansion techniques, and it is capable of performing high throughput 3D multicolor imaging of centimeter scale nervous tissues with micron to sub-hundred nanometer scale spatial resolutions. Nevertheless, we encountered more problems in our effort to establish a method for imaging mouse nervous tissues and decoding large neural networks using our TLS microscope combined with tissue clearing and tissue expansion techniques. First, although tissue expansion enables 3D fluorescence imaging of nervous tissues with the synaptic level spatial resolution. The endogenous fluorophore (EF) preservation efficiency of expanded tissues drops quickly as the tissue size increases, presumably caused by either the protease digestion or the high temperature denaturation and dissociation operations included in the previous tissue expansion methods (51–54). As a result, the EFs are often poorly preserved after expansion for cubic centimeter sized tissues, which makes it necessary to label the expanded tissues via immunostaining after expansion. The problem not only increases the time, cost and difficulty for sample preparation tremendously, but also severely hinders the use of tissue expansion techniques for studying large neural networks at the synaptic level in genetically labeled fluorescent organisms. Second, tissues become more plastic, and the tissue mechanical strength is greatly reduced after clearing and expansion, which make the cleared and expanded tissues extremely fragile, thus difficult to handle and image. For example, the sample could be easily deformed even broken by sample mounting and sample scanning in 3D imaging. This not only makes it difficult to mount the tissue sample on a microscope and conduct the 3D imaging, but also introduces imaging artifacts, reduces the imaging resolution and prevents accurate registration and merging of the subdivided sample volumes for the 3D image of the whole sample. Likewise, the problem becomes more challenging as the tissue size and weight increases. Third, high resolution 3D imaging of large nerves tissues requires not only the ability to image large tissues with high resolution and throughput, but also the ability to process, visualize and manage large image data of at least hundreds of terabytes quickly and efficiently to make the combination of tissue clearing, tissue expansion and LSM techniques practical for decoding large neural networks.

Motivated by the objective to establish a method for imaging large nervous tissues and decoding large neural networks, and to explore the potential of tiling light sheet microscopy combined with tissue clearing and tissue expansion techniques for providing a solution. We developed a comprehensive method that consists of a tissue preparation method and an integrated tissue imaging system to image large genetically labeled nervous tissues from the cellular to synaptic level with high throughput. Specifically, we developed a combined tissue clearing and expansion method to clear and expand mouse CNS tissues at various magnification ratios, preserve the EFs efficiently and maintain the tissue structure and mechanical strength through the sample preparation and imaging processes. We developed an integrated imaging system and workflow for high throughput tissue imaging, image processing and image visualization in parallel. We use our method to study the mouse spinal cord locomotor neural network (SCLNN). We show our method enables high throughput multicolor 3D imaging of large nervous tissues from the cellular to synaptic level and it could advance the decoding of large neural networks significantly.

## Results

### Tissue preparation and parallel tissue imaging, image analysis and image visualization

First, we suspected that the loss of EFs and the low mechanical strength of the expanded tissues are mainly caused by the protease digestion and the high temperature denaturation and dissociation operations included in previous methods (51–54). Since most protein molecules used to label neurons in transgenic animals don’t aggerate, we hypothesized that both protein digestion and dissociation operations can be eliminated to better preserve the EFs without affecting the observation of neuronal and neural network structures. Also enlightened by the findings reported by Susaki et. al. (48) that delipidated tissues have similar properties to electrolyte gels, we hypothesized that the hydrogel monomer permeation and binding efficiency could be higher, and the mechanical strength and structure of the polymerized and expanded tissues can be better preserved by performing dilipidation before the monomer incubation and polymerization instead of after. Thus, we developed a Clearing and Magnification Analysis of Proteome (CMAP) method for large tissue clearing and expansion by modifying the original CUBIC and MAP methods (Fig. 1A). Briefly, CMAP performs tissue dilipidation using CUBIC or similar hydrophilic methods at first, followed by the hydrogel monomer incubation, polymerization, tissue embedding and tissue expansion in sequence. In addition, we developed a low-temperature ultraviolet light induced polymerization method to shorten the hydrogel polymerization process to a few minutes and reduce the influence of the heat generated during polymerization. With the rapid polymerization method, we further developed a layer-by-layer polymerization and embedding procedure to embed the tissue in the same polymer that expands at the same ratio as the tissue during expansion, so that the expanded and embedded tissue is protected and supported mechanically by the hydrogel to better maintain the tissue structure (Fig. 1B and 1C). As expected, tissues prepared with CMAP not only have higher EF preservation efficiency and mechanical strength than that prepared with the original MAP (Fig. 1D, 1E, S1A, and S1B), but also keep the neuronal structures faithfully after tissue expansion (Fig. S1C-S1F), thus allowing direct 3D imaging of genetically labeled mouse CNS tissues after expansion without immunostaining. Furthermore, the expansion ratio with CMAP can be adjusted from 3 to 4 by adjusting the percent composition of the monomer solution compounds (Fig. S1G) (53). As the tissue clearing also introduces an expansion ratio of ∼1.2, the overall tissue expansion ratio with CMAP can be adjusted from 1.2 to 4. It can be chosen according to the neuronal structure to be imaged, the desired spatial resolution and throughput, the tissue size and brightness and the imaging ability of the microscope. This gives a great flexibility in optimizing experiments.

**Figure 1.**
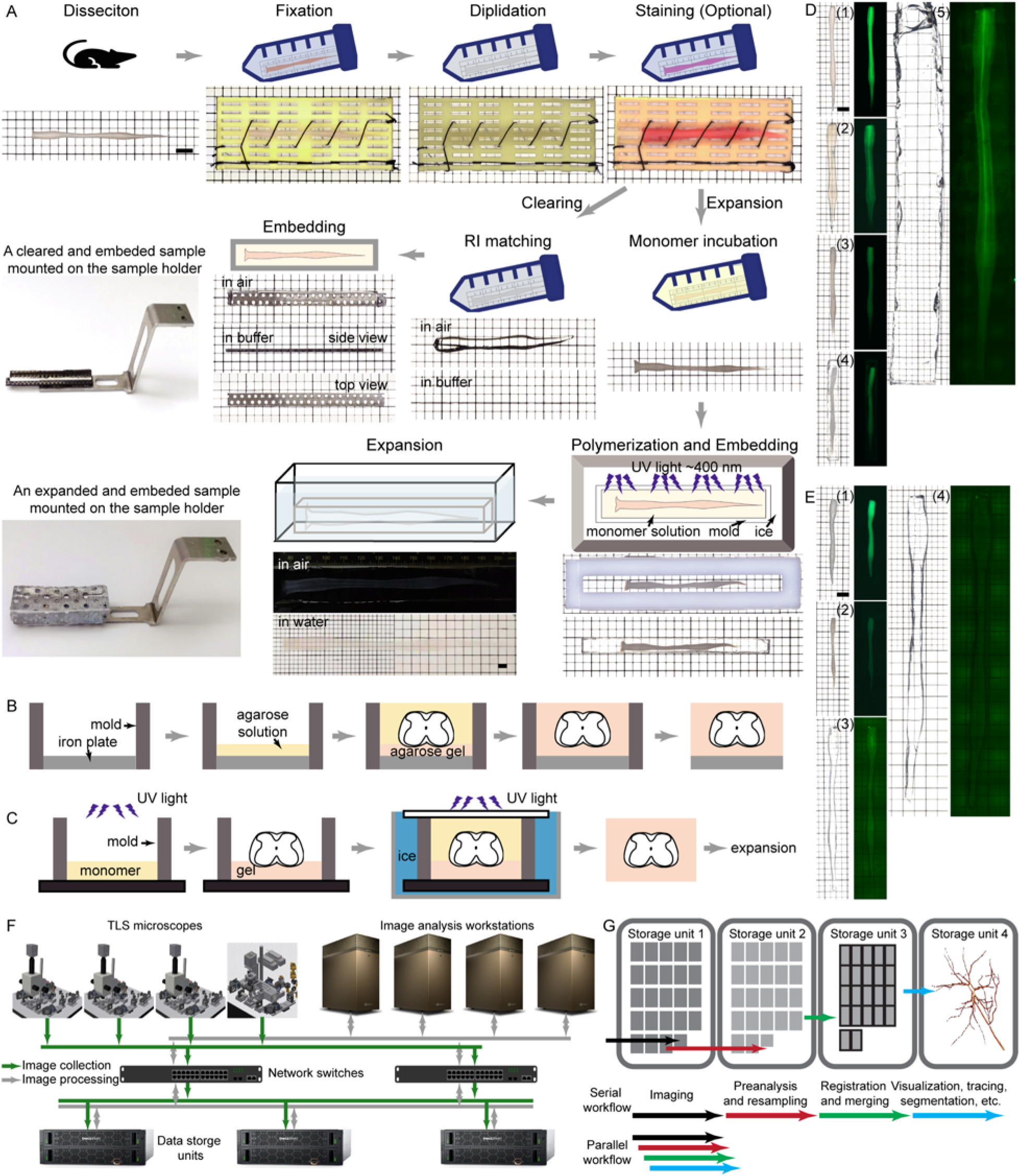
Sample preparation with CMAP, and the parallel tissue imaging, image analysis and image visualization with the integrated imaging system. (A) Tissue clearing and tissue expansion procedure with CMAP. (B) The embedding procedure for cleared tissues. (C) The polymerization and embedding procedure for expanded tissues. (D, E) The comparison of adult Thy1-eGFP mouse spinal cords expanded using CMAP (D): (1) fixation, (2) delipidation, (3) monomer incubation, (4) polymerization and embedding, (5) expansion, and MAP (E): (1) fixation and monomer perfusion, (2) hybridization and polymerization, (3) denaturization and dissociation, (4) expansion, showing the improved EF preservation efficiency. (F) The configuration of our integrated imaging system that consists of a tissue imaging cluster, a high speed data storge cluster and an image processing cluster. (G) The improved throughput with the parallel tissue imaging, image processing and visualization workflow. Scale bars, 5 mm.

Next, we constructed an integrated tissue imaging system that consists of a microscope cluster, a high-speed data storage cluster and an image analysis cluster for tissue imaging, image processing and image visualization in parallel (Fig. 1F and 1G). The microscope cluster includes four TLS microscopes of two configurations (Fig. S2), in which three of them are identical to what we reported previously (65, 66). These microscopes conduct multicolor imaging sequentially, so that the total imaging time elongates proportionally to the number of color channels. It becomes more of a problem as the tissue size and spatial resolution increases. Therefore, we constructed the fourth TLS microscope to perform simultaneous dual-color imaging. Two independently controlled tiling light sheets with excitation wavelengths of 488 nm and 561 nm illuminate the sample simultaneously during imaging, and the corresponding fluorescence signals are imaged to two synchronized cameras at the same time through the same detection objective for simultaneous dual-color imaging. The four microscopes have five sCMOS detection cameras in total resulting in a ∼5 GB/s maximal data collection rate. We built an expandable high speed data storage cluster including multiple data storage units and network switches to fulfill the demand for the data read/write bandwidth and storage space. The cluster provides a ∼17 GB/s read/write bandwidth and a ∼6 PB storage space divided into nine ∼700 TB sectors to save raw and processed images separately. The image analysis cluster consists of multiple image analysis workstations for image processing and visualization in parallel. Each workstation used for microscope control and image analysis is connected to the data storage cluster through an ethernet adapter and an optical cable providing a ∼3 GB/s bandwidth. With the imaging system, a sample is imaged with one of the microscopes by imaging a series of subdivided sample volumes sequentially (65). The raw images of each subvolume are written in a raw 3D image file and processed as soon as the imaging of the subvolume is finished (Fig. 1G). The image registration and merging of the imaged subvolumes are performed along with the imaging process. The parallel imaging, image processing and image visualization workflow allows continuous monitoring and necessary interfering of the experiment, so that the 3D image of the sample is obtained almost instantly after imaging. The overall throughput of our imaging system can be further increased by adding additional units in each cluster. Thus, we obtained the ability and throughput to image large nervous tissues of genetically labelled fluorescent mice from the cellular to synaptic level with CMAP and our integrated imaging system.

### Imaging the mouse spinal cord from the cellular to synaptic level

The spinal cord is primarily responsible for the sensation and locomotion of mammals to interact with its surroundings. Even the most basic movements require precise coordination of many somatic muscles innervated by the spinal cord motor neurons (SpMN), which is also the output layer of the SCLNN. Although significant advances have been made in understanding the SCLNN, and it is possible to reveal the basic role of certain neuron populations in locomotion by studying genetically modified model organisms using electrophysiological, optogenetic, EM and FM techniques (68–72). The mechanism study, even for the most basic locomotor functions remains challenging due to the lack of the ability to observe the corresponding neural circuits with necessary dimension scales and structural details. We decided to focus our study on the motor neuron layer of the mouse SCLNN to explore the utility of our method for decoding large neural networks.

We verified the ability of our method for imaging the mouse SCLNN with the cellular to synaptic level structural details. We first imaged a cleared adult Thy1-eGFP mouse spinal cord from the cervical to the sacral segment, which includes 455 0.65×0.65×4 mm^3^ subvolumes, at a spatial resolution of ∼0.56×0.56×1.5 µm^3^ in ∼100 hours (Fig. 2A, Video S1). As the spinal cord was expanded by ∼1.2 times in each dimension after clearing, the real resolution is ∼0.46×0.46×1.2 µm^3^ according to the original tissue size. Both the spatial distribution, dendritic morphology and axonal projection of the labeled neurons and various spinal tracts allowing the information exchange between different neural layers can be observed clearly (Fig. 2B-2K). For example, the corticospinal tract (CST) axons originating from the cerebral cortex split to multiple axon bundles and decussate at the pyramidal decussation, and the crossed axons rejoin and descend along the dorsal column of the spinal cord after the decussation. The axons originating from the brain stem motor neurons descend along the lateral and ventral funiculi of the spinal cord and develop rich collateral branches sprouting at nodes of Ranvier to spread the information to different spinal cord sections. These axons are much thicker than the CST axons indicating higher information transmission bandwidth and more active activities of these axons (73). The afferent axons of the dorsal root ganglion (DRG) sensory neurons bifurcate to two tracts transmitting in opposite directions along the dorsal funiculus, in which the ascending tract terminates in the brain stem and passes the information to the secondary neurons in the region. Meanwhile, both tracts also develop abundant collateral branches to spread the environmental information to various spinal cord sections.

**Figure 2.**
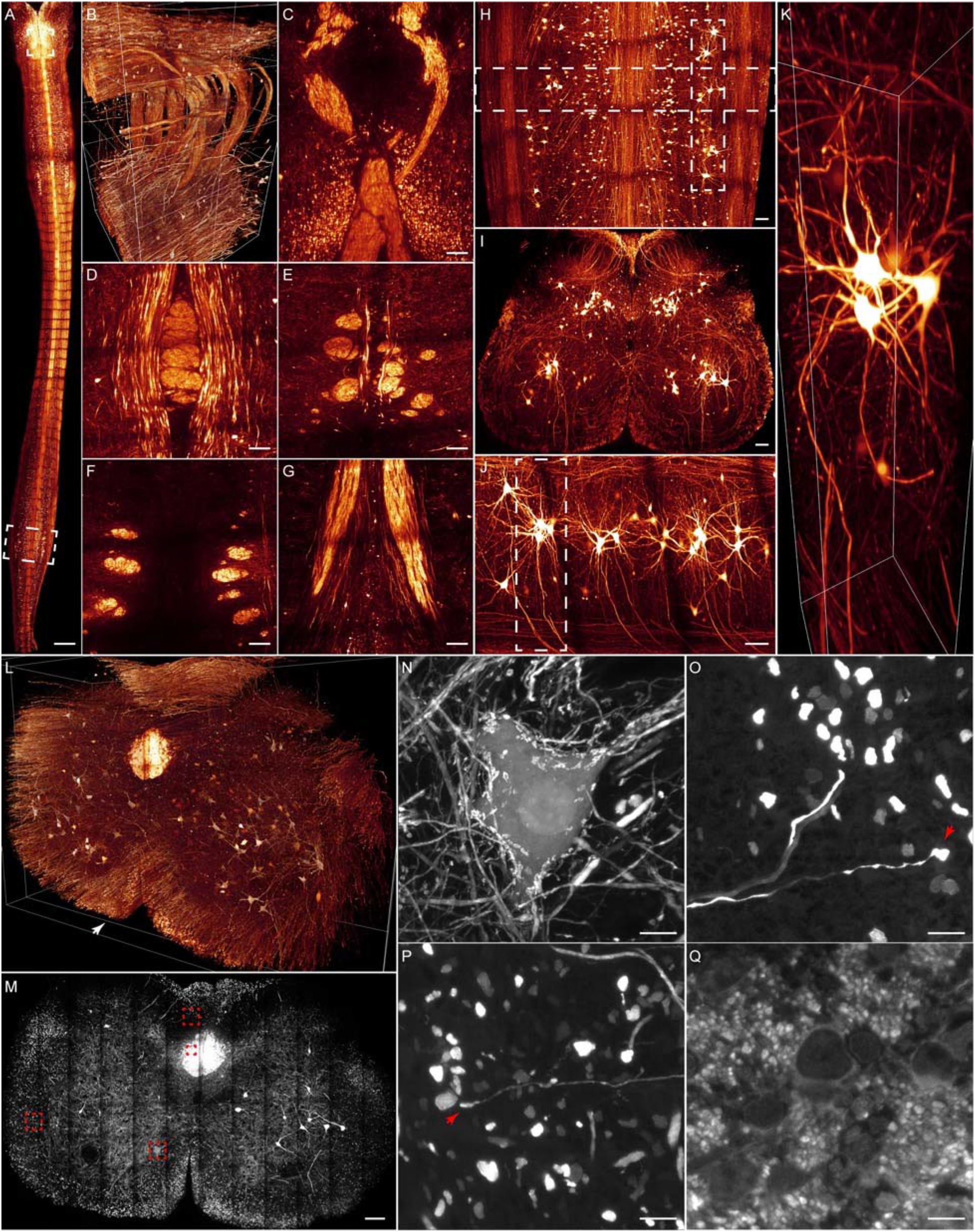
3D imaging of the mouse spinal cord with the cellular to synaptic level spatial resolutions. (A) 3D volume rendering of a cleared Thy1-eGFP mouse spinal cord. (B) 3D volume rendering of the pyramidal decussation indicated in (A). (C-G) Axial and lateral slices through the indicated planes in (B). (H) Lateral maximum intensity projection (MIP) of the selected lumbar section in (A). (I, J) Axial MIPs of the selected regions in (H). (K) 3D volume rendering of the selected region in (J). (L) 3D volume rendering of an expanded adult Thy1-eGFP mouse cervical spinal cord section. (M) Axial MIP of a ∼8 μm thick slice through the indicated plane in (L). (N-Q) Zoom-in views of the selected regions in (M) showing the structure of synaptic connections, axon collateral branches and closely packed CST axons. Scale bars, 1 mm (A), 100 μm (C-G, M), 10 μm (N-P), 5 μm (Q).

We further imaged an expanded cervical spinal cord section of an adult Thy1-eGFP mouse (Fig. 2L-2Q, Video S2), which includes 96 0.6×0.6×3 mm^3^ subvolumes, with ∼0.5×0.5×0.8 µm^3^ spatial resolution in ∼24 hours. The actual spatial resolution is ∼120×120×200 nm^3^ due to the ∼4 times expansion ratio. The result allows the observation of the mouse SCLNN with much more structural details, such as the dense synapses spreading on the spinal cord neurons, the numerous collateral branches of the descending and ascending axons, the spiral projection trajectories of the closely packed CST axons and the complicated connections of the spinal cord neurons. The results confirm our ability to acquire the structural information needed to decode a neural network that includes the spatial organization, dendritic morphology, axonal projection and synaptic connection of neurons. It is worth mentioning that although the highest spatial resolutions of our imaging system are ∼70×70×100 nm^3^ and ∼260×260×400 nm^3^ with and without the ∼4 times tissue expansion (65), we mostly choose the spatial resolutions that are just sufficient to resolve the desired neuronal features for higher throughputs and less photobleaching of samples.

### The organization and morphology of spinal cord motor neurons

Locomotion is mostly realized through the coordination of numerous motor units commanded by the SpMNs. The SpMNs arrange into various motor columns to operate (74–76). However, the organization of the SpMNs and motor columns in mice hasn’t been visualized with sufficient detail (77, 78). As all SpMNs are cholinergic, we imaged a cleared adult ChAT-eGFP mouse spinal cord from the cervical to the sacral segment to gain more details (Fig. 3A, Video S3 and S4). We segregated the spatially distinct motor neurons in the lateral and ventral horns, and we identified eleven pairs of bilateral columns (Fig. 3B-3F) and two unilateral columns. In which the bilateral columns consist of six cervical pairs (Ca-Cf), one thoracic pair (T), three lumbar pairs (La-Lc) and one pair through the whole spinal cord (MMC), and the two unilateral columns locate at the boundaries of the thoracic, lumbar and sacral segments respectively (Ma, Mb). We counted the SpMNs in all columns and identified 29,392 SpMNs (Fig. 3G). The result suggests the motor neuron layer of the mouse SCLNN consists of a little more than 30K motor neurons, and the locomotion of mice is realized through the coordination of these neurons. The control diagram of them would need to be depicted to understand the mouse SCLNN. Meanwhile, the slightly uneven distribution of the motor neurons on both sides of the spinal cord could result in asymmetrical locomotor abilities, and the larger numbers of motor neurons and motor columns in the cervical segment than that in the lumbar segment suggests finer locomotion control ability of the forelimbs than hindlimbs in mice.

**Figure 3.**
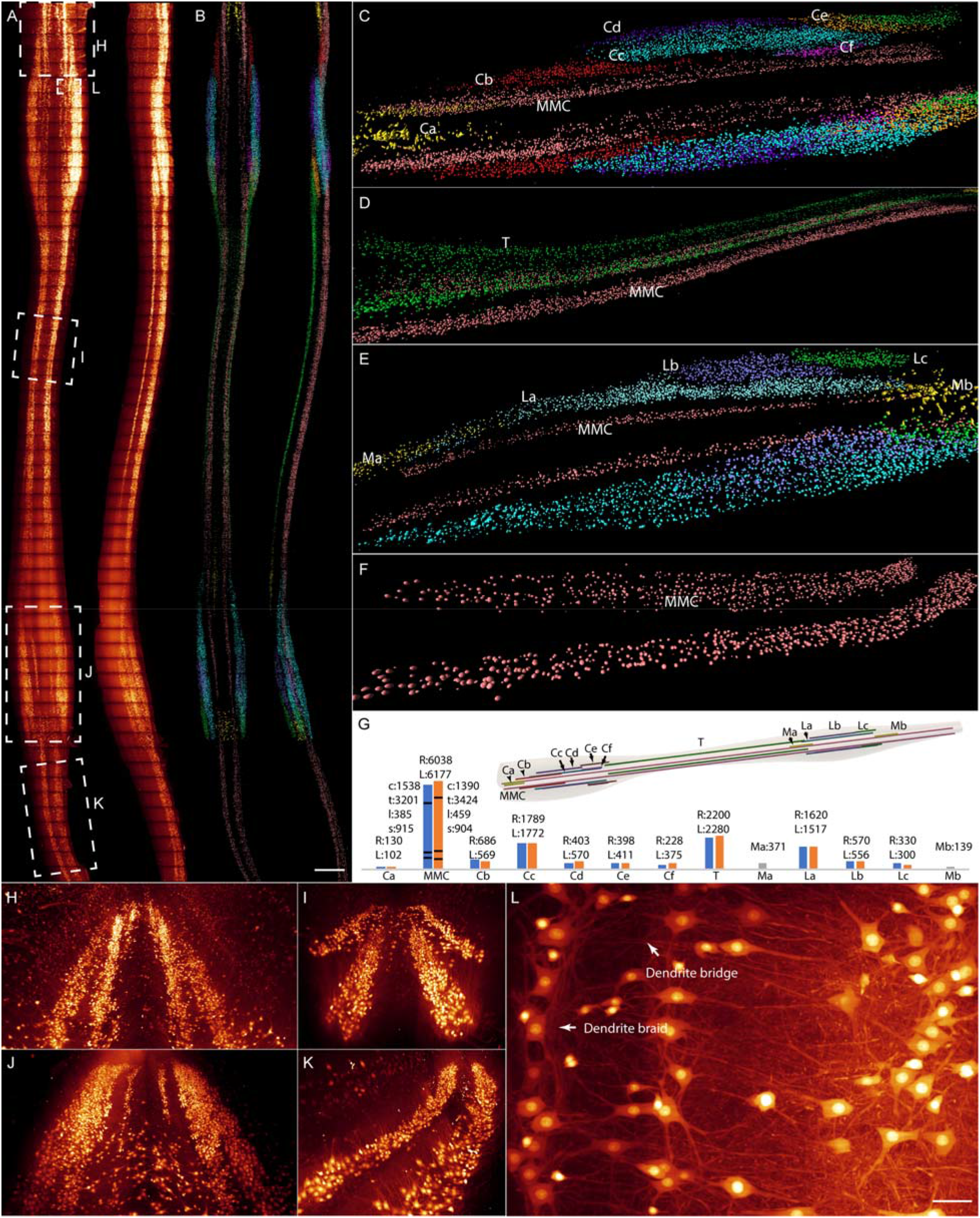
The organization of the SpMNs in mouse spinal cord. (A) Lateral and axial MIPs of a cleared adult ChAT-eGFP mouse spinal cord. (B) The segregation of the SpMNs and moto columns. (C-F) Zoom-in views of the cervical, thoracic, lumbar and sacral spinal cord sections showing the spatial segregation of various motor columns. (G) The number of SpMNs in each identified motor column. (H-K) 3D volume renderings of the selected regions in (A) showing the dendritic territory and morphology of different motor columns. (L) Lateral MIP of a 100 μm thick ventral horn region indicated in (A) showing the dendrite braid and bridge structures of the SpMNs. Scale bars, 1 mm (B), 50 μm (L).

The dendritic span and morphology of a motor neuron determine its ability and location to receive signals from the upper neural layers and are therefore critical for the operation of the SCLNN. Our result makes the information easily accessible. It shows that the dendrites of the motor neurons in different columns have both exclusively occupied territories and spaces shared with other columns (Fig. 3H-3K and S3). For example, only the medial column motor neurons have dendrites growing towards the spinal cord midline, and only the lateral column motor neurons have dendrites growing towards the ipsilateral lateral horn. While both medial and lateral column motor neurons have dendrites extending to each other. The morphologies suggest the SpMNs in different columns could both receive unique information to innervate specific muscles and share common information to coordinate intermuscular motor activities. We also found that many SpMNs have dendrites growing both along the spinal cord rostrocaudal axis, forming intracolumn braid-like structures, and in the transverse directions toward the neighbor columns, forming intercolumn bridge-like structures (Fig. 3L). Presumably, these structures could facilitate the sharing of the input projections from upper neural layers to coordinate intramuscular and intermuscular behaviors. Particularly, the braid structure allows an input axon to summon more intracolumn motor neurons from near to far as the axon signal becomes stronger regardless of the projection position of the axon to the motor column. It could be a complementary mechanism of the Henneman size principle (79–82) to generate gradient forces as it relaxes the requirement for an axon to project to specific motor neurons of various sizes. Similarly, the dendrite bridges that cross adjacent motor columns could make it easier for an input axon to coordinate intercolumn motor activities by projecting to these dendrite bridges.

To better observe the dendritic morphology of individual SpMNs, we imaged a cleared adult wild-type (WT) mouse spinal cord with the motor neurons sparsely labeled through muscle injection of rAAV2-retro-hSyn-tdTomato vectors (83), and we segmented thirty SpMNs from different locations of the spinal cord (Fig. 4A-4K, Video S5). Despite the distinct morphologies of the segmented motor neurons, there are many similarities. Most motor neurons have a 1-2 mm dendrite span and 4-8 primary dendrites that grow mostly along the rostrocaudal and transverse directions, which is consistent with the dendrite braid and bridge structures observed in ChAT-eGFP mice. The motor neurons with fewer primary dendrites are more polarized and tend to grow in transverse directions. A small portion of motor neurons develop 1-2 axon collateral branches indicating the participation of these neurons in recurrent circuits (Fig. 4K) (84). It is critical to identify the specific muscles and motor activities for which these motor neurons are responsible in order to understand the function of recurrent circuits in locomotion control. Surprisingly, some motor neurons have the axon and dendrites grow towards the opposite sides of the spinal cord. Further study is required to verify the function of these neurons.

**Figure 4.**
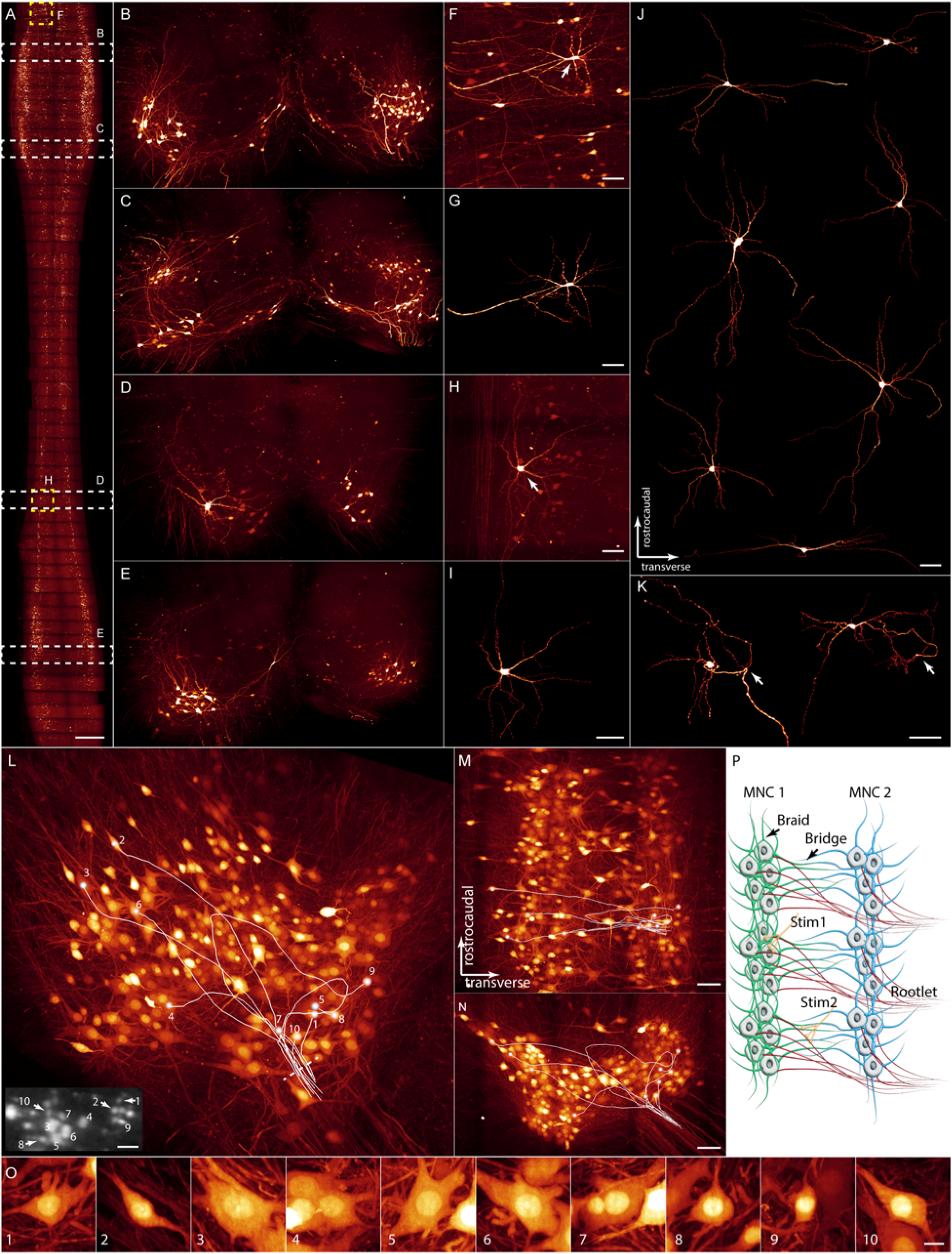
The dendritic morphology of individual SpMNs, and the structure of spinal cord ventral rootlets. (A) Lateral MIP of a cleared adult WT mouse spinal cord with the SpMNs sparsely labeled. (B-E) 3D volume renderings of the selected spinal cord sections in (A) showing the dendritic morphology of individual SpMNs in various motor columns. (F-I) Zoom-in views of the selected regions in (A) and the segmented motor neurons. (J) Lateral MIPs of multiple segmented SpMNs. (K) SpMNs with multiple axon collateral branches. (L-N) 3D volume rendering and MIPs of an expanded adult ChAT-eGFP mouse cervical spinal cord ventral horn section showing the distribution of the SpMNs with axons in the same rootlet. The insert in (L) shows the cross section of the rootlet at the indicated position in (L). (O) Zoom-in views of the traced motor neurons in (L). (P) A proposed model describing the organization of the SpMNs for receiving and sending signals. Scale bars, 1 mm (A), 100 μm (F-K), 5 μm (L insert), 200 μm (M, N), 10 μm (O).

The efferent axons of the SpMNs combine to a series of ventral rootlets before leaving the spinal cord. Despite the discovery of the structure long ago (71), the distribution of the motor neurons with the axons joining in the same rootlet is unclear. To answer the question, we imaged the cervical, thoracic and lumbar spinal cord sections of an expanded adult ChAT-eGFP mouse spinal cord, and we traced the axons in the same rootlet in each section (Fig. 4L-4O and S4, Video S3 and S6). We found that the SpMNs with axons in the same rootlet stay closely along the rostrocaudal axis despite being located in different ipsilateral columns, except in the thoracic segment where all SpMNs locate in the MMC column. The formation of rootlets seems to be natural, as the efferent axons take the shortest path to leave the spinal cord. The structure allows the neural impulses fired from the related SpMNs in adjacent spinal sections and motor columns to travel nearly the same physical path to reach the innervated muscle fibers, thus ensuring accurate intermuscular coordination.

As shown, in addition to a better understanding on the distribution and morphology of the SpMNs, our results suggest a model that the SpMNs develop intracolumn dendrite braids and intercolumn dendrite bridges, and the SpMN efferent axons form rootlets to better coordinate the intramuscular and intermuscular motor activities (Fig. 4P). We further imaged the cleared spinal cords and expanded spinal cord sections of several juvenile ChAT-eGFP mice (P1, P7, P28) (Fig. S5, Video S4), and we noticed significant changes in the quantity, distribution and morphology of the SpMNs with the growth of mice. Apparently, our method can also be used to study the development and learning processes of the mouse SCLNN which have been nearly impossible to conduct due to the lack of tools to image large nervous tissues with sufficient spatial resolution and throughput.

### The projection and connection of multiple spinal tracts for locomotion control

The functioning of the SCLNN relies on the information conveyed to the spinal cord through various axon tracts whose projection patterns determine how and where the carried information is distributed. In the mouse SCLNN, the descending axon tracts originating from the cerebral cortex and brainstem deliver high level movement commands to drive the SpMNs directly or indirectly. While the afferent axons of the dorsal root ganglion (DRG) sensory neurons not only bring in the exteroceptive and proprioceptive sensation information to the CNS but also mediate numerous reflex movements. As these axon tracts initiate different types of locomotion, such as the reflexive, rhythmic and voluntary movements. We wondered whether and how the projection pattern of these spinal tracts establishes the structural foundation of their roles in locomotion control.

We first studied the axonal projection of the sensory neurons in the mouse spinal cord. We imaged a cleared WT mouse spinal cord with the right L4 DRG sensory neurons sparsely labeled through the injection of rAAV9-hSyn-Cre and rAAV9-CAG-DIO-tdTomato vectors into the DRG (Fig. 5A). The afferent axons of the labeled neurons divide into the ascending and descending tract after entering the spinal cord. Both tracts develop rich collateral branches within a limited distance from the entry location of the afferent axons. Most of the branches terminate in the ipsilateral dorsal horn, a small number of them reach the ipsilateral ventral horn and almost no branches cross the spinal cord midline (Fig. 5B-5G). To better visualize the axonal projection of individual sensory neurons, we imaged another cleared WT mouse spinal cord with the sensory neurons sparsely labeled through the muscle injection of rAAV9-hSyn-Cre and rAAV9-CAG-DIO-tdTomato vectors (85), and we traced the afferent axons and primary collateral branches of several sensory neurons with the neuron soma in different DRGs (Fig. 5K-5O, Video S7). We found that both the ascending and descending axons develop several collateral branches only within roughly a centimeter range around the afferent axon entry position with most collateral branches project exclusively to the ipsilateral dorsal horn of the spinal cord. The descending axons terminate after projecting for a few millimeters, and the ascending axons merge into the dorsal fasciculus and keep ascending without further branching until terminating in the brainstem. Both results are consistent and suggest that the primary function of sensory neurons is to bring the sensation information to the brain for further processing and to the spinal cord local circuits within a limited range to trigger reactions to stimuli, but only a small number of sensory neurons can interact with the SpMNs directly to initiate movements. We confirmed the existence of monosynaptic connections between the proprioceptive sensory neurons and SpMNs by imaging an expanded lumbar spinal cord section of an adult ChAT-eGFP mouse with the right L1 DRG proprioceptive neurons sparsely labeled through the injection of rAAV9-PV-Cre and rAAV9-CAG-DIO-tdTomato vectors into the DRG (Fig. 5P-5Y, Video S8). As expected, although most axon collateral branches of the proprioceptive neurons still terminate in the ipsilateral dorsal horn, a significant number of them terminate in the ipsilateral ventral horn and form direct connections with the SpMNs. It can also be observed that the axon branches growing in both rostrocaudal and transverse directions to form synapses with intracolumn and intercolumn SpMNs (Fig. 5R, Video S8). Although the monosynaptic connections between the proprioceptive neurons and SpMNs are unable to initiate complicated locomotion. The simple yet robust connections ensure the faithful and efficient execution of the most basic reflex movements in mice.

**Figure 5.**
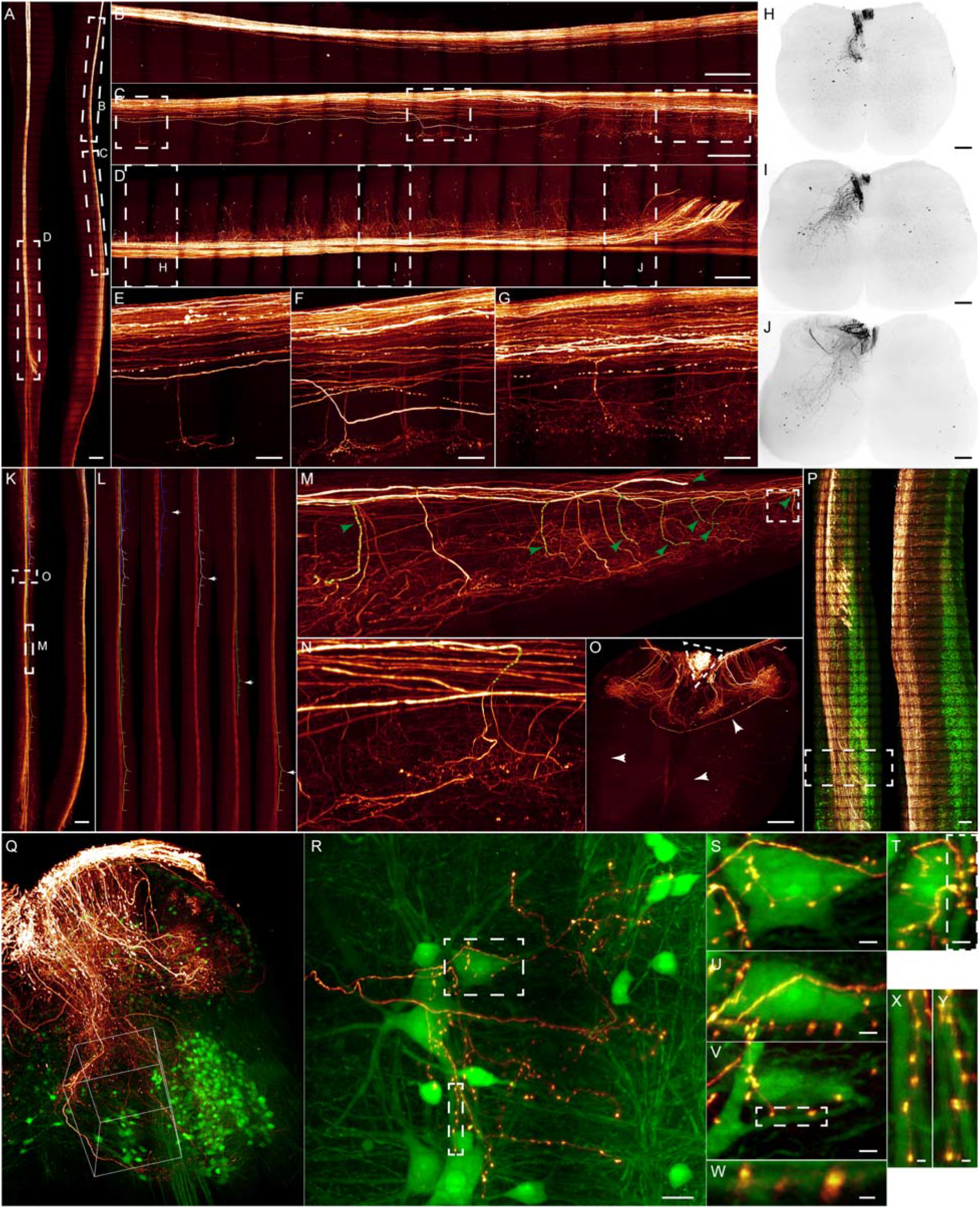
The afferent axonal projection of sensory neurons in mouse spinal cord. (A) Lateral and axial MIPs of a cleared adult WT mouse spinal cord with the right L4 DRG sensory neurons sparsely labeled. (B-D) Zoom-in views of the selected regions in (A) showing the different numbers of axon branches in various spinal cord sections. (E-G) Zoom-in views of the selected regions in (C) showing the axon branch morphology. (H-J) Axial MIPs of the selected regions in (D) showing the projected regions of the axon branches in various spinal cord sections. (K) Lateral and axial MIPs of a cleared adult WT mouse spinal cord with sensory neurons sparsely labeled. (L) The projection of the sensory neuron afferent axons entering the spinal cord at various spinal cord sections. (M) 3D volume rending of the selected region in (K). (N) Zoom-in view of the indicated axon terminal arbor in (M). (O) Axial MIP of the selected section in (K) showing the projected areas of the sensory neuron axon branches. (P) Lateral and axial MIPs of an expanded ChAT-eGFP mouse lumbar spinal cord section with the right L1 DRG proprioceptive sensory neurons sparely labeled. (Q) 3D volume rending of the selected region in (P). (R) Lateral MIP of the indicated volume in (Q). (S-U) Lateral MIPs of the indicated region in (R). (V) Lateral MIP of the selected region in (T) showing the monosynaptic connections between the proprioceptive sensory neurons and motor neurons. (W) Axial MIP of the selected regions in (V). (X, Y) Lateral and axial MIPs of the selected region in (R). Scale bars, 1 mm (A,K, and P), 500 μm (B-D), 100 μm (E-G), 200 μm (H-J,O,P), 20 μm (R), 5 μm (S-V), 2 μm (W-Y).

The upper motor neurons in the cerebral motor cortex and brainstem are essential to initiate voluntary, rhythmic and more complicated reflex movements. In which the motor cortex is more responsible for skilled movements, and the brainstem is more effective in orchestrating basic movements, such as balance maintenance and posture control (86). We imaged several spinal tracts originating from the motor cortex and brainstem and compared their axonal projection patterns to understand the influence of the axonal projection patterns on the roles the various neuron populations in mouse locomotion control.

The cerebral cortex motor neurons arrange somatotopically to manipulate the movements of different body parts in mammals (87–90). To understand how well the point-for-point correspondence relationship is established in mice, we studied the axonal projection of the upper motor neurons in the motor cortex that evoke forelimbs (FL) and hindlimbs (HL) respectively. We imaged the cleared spinal cords of two adult WT mice injected with rAAV2-cFos-FLPo and rAAV2-CAG-fDIO-eYFP vectors into the right FL and HL regions respectively (Fig. 6). The results show that both the FL-spinal tract and the HL-spinal tract project along the CST but terminate in the cervical and lumbosacral spinal cord sections respectively. Abundant axon branches were observed in both tracts. Most FL-spinal tract axon branches project to the ipsilateral lateral and ventral horns, and a small number of branches project to the contralateral lateral and ventral horns either directly or by crossing the midline from the ipsilateral side of the spinal cord (Fig. 6C-6E, Video S9). In contrast, the HL-spinal tract axon branches mostly project to the ipsilateral dorsal horn in the cervical and upper thoracic sections and project to the ipsilateral dorsal, lateral and ventral horns in the lumbosacral sections (Fig. 6F-6I, Video S10). A small number of branches also terminate in the contralateral side of the spinal cord. We repeated the experiment with lower neuron labeling densities, and we traced multiple axons in both tracts to inspect the projection pattern of individual CST axons (Fig. S6). As shown, the FL-spinal tract axons terminate and develop multiple collateral branches in various cervical sections, and the HL-spinal tract axons could have both bifurcations and collateral branches along the entire spinal cord with most branches concentrated in the lower thoracic and lumbosacral sections. In consequence, the FL-spinal tract delivers the FL neural pulses to the spinal cord cervical sections exclusively while the HL spinal tract could deliver the HL neural pulses to almost the whole spinal cord besides the lumbosacral sections that are responsible for hindlimb control. The results also suggest that the mouse motor cortex neurons only establish a rough point-to-point correspondence relationship with the SpMNs through the CST, and the somatotopic relationship varies for different cortex regions and body parts. This could lead to different fine control ability of forelimbs and hindlimbs in mice.

**Figure 6.**
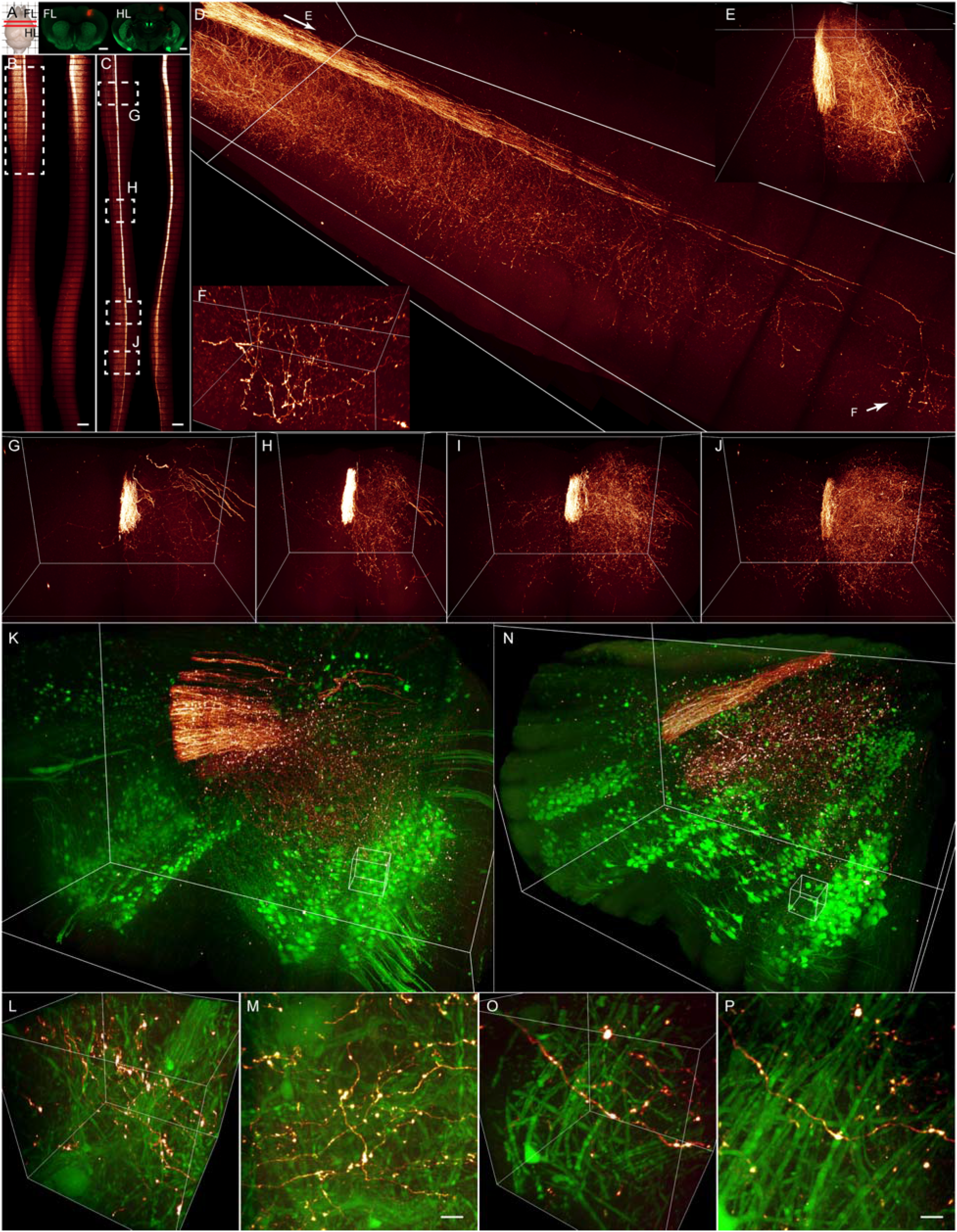
The CST projection in mouse spinal cord. (A) Spare labeling of the FL-spinal and HL-spinal tract via motor cortex virus injection. (B, C) Lateral MIPs of two cleared WT mouse spinal cords showing the FL-spinal tract and HL-spinal tract projection. (D) 3D volume rendering of the selected section in (B). (E) Prospective view of the section from the indicated direction in (D). (F) Zoom-in view of the indicated axon terminal arbor in (D). (G-J) 3D volume rendering of the selected sections in (C) showing the HL-spinal tract axon branch projection in various spinal sections. (K) 3D volume rendering of an expanded spinal cord cervical section of an adult ChAT-eGFP mouse with the FL-spinal tract sparsely labeled. (L, M) 3D volume rendering and lateral MIP of the selected volume in (J) showing no monosynaptic connections. (N) 3D volume rendering of an expanded spinal cord lumbar section of an adult ChAT-eGFP mouse with the HL-spinal tract sparsely labeled. (O, P) 3D volume rendering and lateral MIP of the indicated volume in (N) showing no monosynaptic connections. Scale bars, 1 mm (A-C), 10 μm (M, P).

Although it is believed that CST plays a critical role in skilled movements, the interaction between the CST and the SpMNs could be direct or indirect in different mammalian species (86, 91). The existence of monosynaptic connections between the CST and SpMNs in mice has been debated for a long time (92–95). A high certainty conclusion is difficult to reach because it requires the visualization of both CST and SpMNs, and also the screening of the mouse spinal cord with synaptic level spatial resolution which is difficult to conduct with conventional techniques. To answer the question, we imaged the expanded cervical and lumbar spinal cord sections of several ChAT-eGFP mice injected with rAAV2-cFos-FLPo and rAAV2-CAG-fDIO-tdTomato vectors into the FL and HL cortex regions respectively (Fig. 6J-6O, Video S11 and S12). We found that very few CST axons reach the territories occupied by motor neurons despite the rich CST axon terminal branches in the ventral horns, and we didn’t find any monosynaptic connections between the CST and SpMNs despite the existence of abundant boutons along the CST axon terminal branches. In addition, both the projection patterns and the high number of unmyelinated axons of the mouse CST don’t seem to favor the accurate and efficient information distribution that is likely needed for fine locomotion control (Hsu et al., 2006). We therefore conclude that the CST doesn’t drive the SpMNs directly in mice. CST is likely to play a different role in mice than that in primates (94), and the mouse probably has a limited ability for skillful movements.

The brainstem is another command center for locomotion control in mice. The upper motor neurons in different brainstem regions participate in various motor activities by projecting their axons to the spinal cord through various descending tracts. In which the rubrospinal tract (RST) originating from the red nucleus (RN) plays a critical role in limb control (97–99), and the reticulospinal tract (ReST) originating from the reticular information (RI) is essential for postural control and balance maintenance (100–105). We investigated the projection patterns of both tracts to seek the structural characteristics that could facilitate their functioning.

We first imaged a cleared WT mouse spinal cord with the RST axons sparsely labeled through the injection of rAAV2-cFos-FLPo and rAAV2-CAG-fDIO-eYFP vectors into the right RN (Fig. 7A). The RST axons descend in the dorsolateral funiculus after decussating at the ventral tegmentum and terminate from the cervical to sacral spinal cord sections. Along the descending path, the RST axons develop rich collateral branches that are predominantly confined in lamina V-VII and X, project medially into the spinal cord gray matter and terminate in the medial regions without crossing the middle line (Fig. 7B-7E). The branches are also more concentrated in the cervical and lumbar sections, which indicates the more active involvement of the RN neurons in limb control of mice. We traced multiple RST axons to examine how information is distributed by individual RST axons (Fig. 7F-7l). As shown, an RST axon could develop numerous collateral branches to deliver the neural pulses to a large range of the spinal cord. More importantly, we identified two groups of axons that project to cervical sections and lumbosacral sections exclusively. This echoes previous findings that RN neurons have a somatotopic organization in relation to their projections to the spinal cord cervical and lumbosacral sections (106, 107), and suggests the RN neurons could enable independent control of each limb in mice through the RST. Our results are also coherent with the observation that RN is almost invariably present in animals having fins, wings, limbs or limb-like structures as a mean of locomotion, and is absent in primitive vertebrates (97). Despite the critical role of the RN neurons in limb control of mice, there is little evidence regarding whether the RN neurons could drive the SpMNs directly through the RST. Küchler et al. reported that the RST axons could drive the distal forelimb muscle motor neurons directly in rats, but the observation was made by imaging rat spinal cord tissue slices using a widefield fluorescence microscope, which hardly offers sufficient spatial resolution and 3D resolving ability (108). To verify the report, we imaged the expanded cervical and lumbar spinal cord sections of several adult ChAT-eGFP mice with rAAV2-cFos-FLPo and rAAV2-CAG-fDIO-tdTomato vectors injected into the right RN. We found that the RST axons branches rarely project to the spinal cord ventral horns, and we didn’t find any monosynaptic connections between the RST axons and the SpMNs. Our result suggests that the RST axons don’t drive the SpMNs directly in mice, which is different from the conclusion made by Küchler et al. in rats.

**Figure 7.**
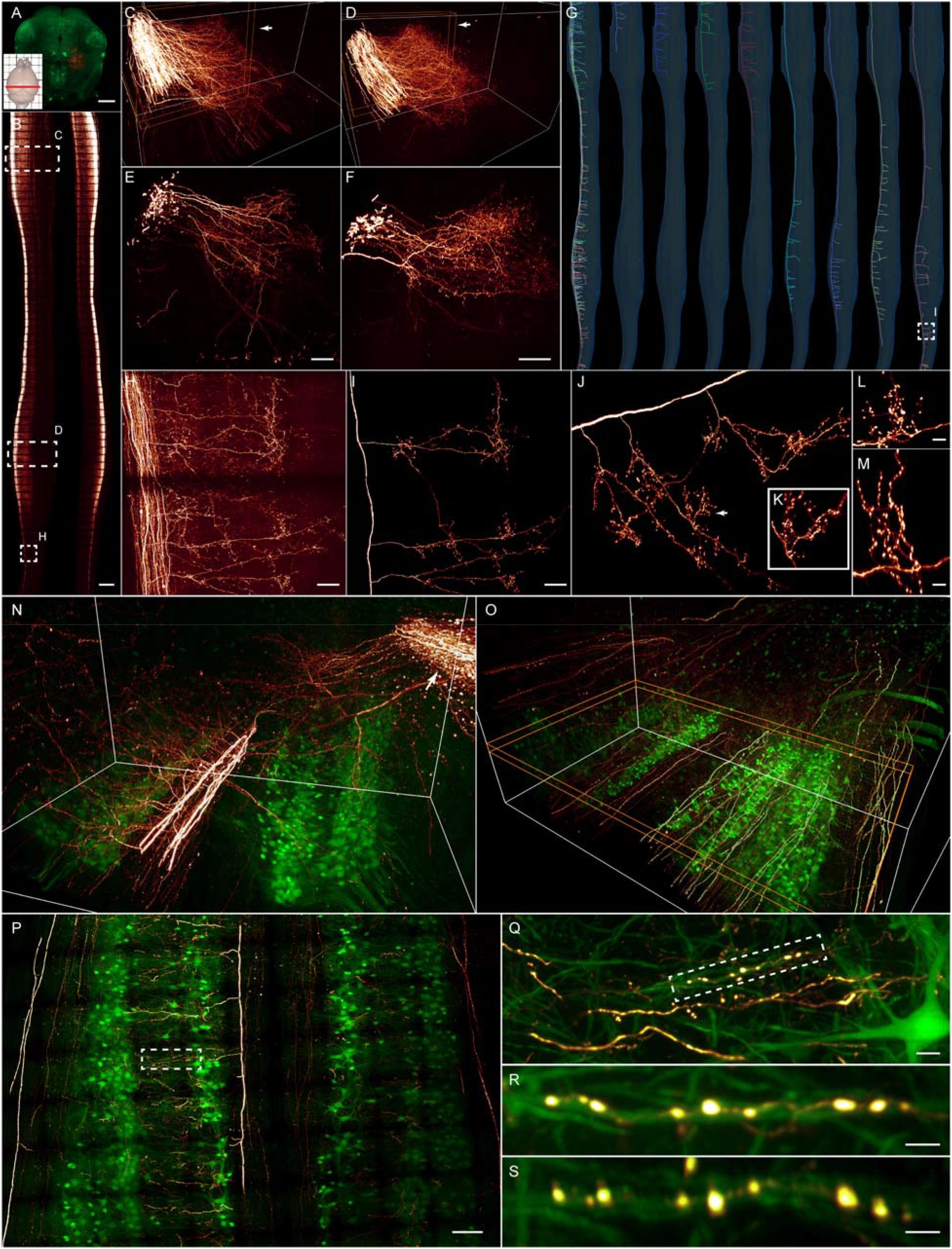
The RST and ReST projections in mouse spinal cord. (A) Spare labeling of the RST via RN virus injection. (B) Lateral and axial MIPs of a cleared adult WT mouse spinal cord with sparsely labeled RST axons. (C, D) 3D volume renderings of the selected cervical and lumbar sections in (B). (E, F) Axial MIPs of the indicated 200 μm thick slices in (C) and (D) showing the projection pattern of the RST axon branches. (G) Traced individual RST axons showing the extensive branches and the somatotopic projection patterns of the RST. (H, I) Zoom-in views of the indicated area in (B) and (G) showing the extracted axon branches. (J) 3D rendering of the extracted axon branches. (K-M) 3D rendering and the lateral and axial MIPs of the indicated terminal branch in (J). (N) 3D volume rendering of an expanded cervical spinal cord section of an adult ChAT-eGFP mouse with the RST sparsely labeled. (O) 3D volume rendering of an expanded cervical spinal cord section of an adult ChAT-eGFP mouse with the ReST sparsely labeled. (P) Lateral MIP of the indicated 50 μm thick slice in (O). (Q) Zoom-in views of the indicated area in (P). (R, S) Lateral and axial MIPs of the selected region in (Q) showing the direct connections between the ReST and SpMNs. Scale bars, 1 mm (A and B), 200 μm (E and F), 100 μm (H, I and P), 20 μm (L and M), 10 μm (Q), 5 μm (R and S).

We studied the ReST in the next by imaging the cleared spinal cords and expanded spinal cord sections of adult WT and Chat-eGFP mice injected with rAAV2-cFos-FLPo and rAAV2-CAG-fDIO-tdTomato vectors into the right gigantocellular reticular formation (Gi). The ReST axons also project to the whole spinal cord and develop rich collateral branches. Most ReST axons descend along the ipsilateral ventral and lateral funiculi, and a subset of axons descend along the contralateral ventral and lateral funiculi (109, 110) (Fig. S7A-S7E, Video S14). The lateral and ventral funiculi axon branches tend to project laterally and vertically respectively, and a substantial number of branches in all funiculi cross the midline and terminate in the contralateral side. We traced multiple axons in different funiculi to gain a better understanding on the projection characteristic of individual ReST axons (Fig. S7F-S7L). The ReST axons show much more complicated projection patterns compared to that of the RST axons. In which a key difference is that individual ReST axons could project to either side of the spinal cord at various spinal sections through collateral branches. For example, the branches of the axon G-5 (Fig. 7G) project vertically to the ipsilateral cervical sections, but laterally across the midline to the contralateral thoracic sections. This suggests the ability of the axon to coordinate intermuscular activities of forelimb and body muscles on opposite body sides, and the ability is likely needed for balance maintenance. Indeed, both posture control and balance maintenance could be extremely complicated as they require the rapid and continuous coordination of various somatic muscles across the whole body, and the mechanical responses of the involved muscles must be highly versatile and dynamic under different conditions. This could lead to the diverse projection patterns of the ReST axons.

Riddle et al. showed that the ReST forms direct connections with the SpMNs in anesthetized monkeys by measuring the neural pulses using electrophysiological methods (111), but the direct observation of the connection has never been achieved. Our results show abundant monosynaptic connections between the ReST and the SpMNs in various motor columns (Fig. 7O-7S, Video S15), and confirm the ability of Gi motor neurons to drive the SpMNs directly through the ReST in mice. The presence and absence of direct connections between the SpMNs with ReST and RST could be either the result or cause of the different types of locomotion that the Gi and RN motor neurons are responsible for. Limb movements mediated by the RN through RST should be either voluntary or rhythmic. In which the participation of the spinal inter neurons are required to orchestrate the SpMNs. The corresponding movements could be more intentional, but less urgent. On the contrary, posture control and balance maintenance are mostly involuntary, and the involved muscles need to respond continuously and rapidly to the received proprioceptive information. The monosynaptic connections between the ReST and the SpMNs could ensure the efficient and reliable executions of the movement commands. Indeed, such an involuntary control manner for balance maintenance actually makes it difficult for one to lose balance unless it is done on purpose. Clearly, the unique projection characteristics of the RST and ReST provide the structural basis for the functioning of the RN and Gi in locomotion control.

## Discussion

EM has been the only technique that can be used to image nervous tissues from the cellular to synaptic level. Unfortunately, the low throughput and the inability to take the advantage of genetic labeling tools limit it from being used for decoding large neural networks efficiently. Thus, the essential goal of our research is to develop a method with high throughput and sufficient spatial resolving ability to be used as a substitute or supplement for EM for imaging large nervous tissues and decoding biological neural networks, thus enabling the understanding of mammalian intelligence with logic. The numerous research progresses achieved by the tissue clearing, tissue expansion and light sheet microscopy techniques suggest their combination as a possible method.

Convinced by the potential of the combination, we developed an integrated method to clear, expand and image genetically labeled large nervous tissues, and we demonstrated its utility by studying the mouse SCLNN. Particularly, we investigated the organization, morphology and development of the SpMNs, the projection of multiple spinal tracks and the existence of direct connections between these spinal tracks with the SpMNs. Our method enables the study with both high spatial resolution and throughput that are unachievable before and brings new insights to the logical architecture of the locomotor neural network. More importantly, our method can presumably be used to study other neural networks from various perspectives, such as the developmental and learning processes of neural networks, and this is extremely important for the understanding of biological intelligence but has been impossible to conduct experimentally before.

The fundamental advantages of the presented method come from three aspects. First, it enables the use of various genetic labeling techniques, which not only makes it much more convenient to study specific neural networks, but also reduces the data analysis complexity as data can be visualized instantly after imaging without requiring extensive data analysis. Second, it provides both the ability and flexibility to acquire the structural information of neural networks from the cellular to synaptic level with the optimal efficiency. Although the highest spatial resolution is still an order of magnitude away from that of the EM, the sub-hundred nanometer level resolution in combined with fluorescent labeling still enables the identification of synaptic connections with high certainty. Third, it has a much lower implementation threshold and a much higher throughput compared to that of the EM, thus allowing much easier implementation in various research. Although there is no doubt that EM will remain as the gold standard to examine the ultrastructure of neural networks, the adoption of the presented method would permit the study of biological neural networks to proceed much faster in many directions that have been severely restrained by the limited ability to observe neural networks efficiently with sufficient structural details. The removal of the obstacles could also accelerate the iterative development of the related neuron labeling, tissue preparation, tissue imaging, image analysis and neural network modeling techniques for decoding biological neural networks.

Despite the promising potential of the presented method, critical improvements are needed in many aspects. First, the challenge of studying large neural networks still increases sharply as the nervous tissue size increases. More specific, stable, bright fluorescent probes, higher fluorescent labeling density and efficiency, higher EF preservation efficiency, transparency, expansion ratio, mechanical strength of the prepared tissue, and microscopes that are more capable and efficient to image large tissues with high 3D spatial resolution are always desired. Second, a fundamental advantage of FM over EM is to label neural networks of interest via genetic labeling. The ability to label specific neurons more precisely both in space and time is critical to further exploit the advantage. For example, more effective anterograde and retrograde transsynaptic labeling methods could facilitate the recognition of specific neural layers in complicated neural networks, and the ability to label certain neurons at the desired moments during various developmental and learning processes of a neural network is necessary to understand the maturation of the network regardless of the imaging ability. Third, data management and analysis become a major bottleneck as the imaging resolution, throughput and tissue size increases. Hopefully, the lowered threshold for imaging nervous tissues by the presented method could have more scientists involved to solve the problem. Forth, despite the advances made in mapping neural networks diagrams, it should be aware that the diagram mapping alone is unlikely to lead to the complete understanding of biological neural networks, because all results are collected from ceased neural networks that are supposed to be highly dynamic and variable. Instead, the research would allow scientists to make better hypotheses, and build more accurate neural network models to be verified in live experiments using other methods, such as the electrophysiological, optogenetic and live imaging techniques. Finally, the understanding of biological neural networks is extremely challenging, and it is becoming more and more resource intensive. It is probably worth thinking more carefully about how neural network diagrams could facilitate the understanding of neural networks and finding a solution to make the use of the experimental resources and the produced data more efficient, so that more scientists can participate in the research and benefit from the latest technology advances and research outputs.

## Methods

### Experimental Model and Subject Details

The collection of mice tissues used in this study was approved by the Institutional Animal Care and Use Committee of the Westlake University (Approval No: 19-035-GL). All mouse lines were kept on C57BL/6J background and were housed in a standard 12:12 light-dark cycle. C57BL/6J was purchased from Shanghai Jihui Laboratory Animal Care Co., Ltd.. Thy1-eGFP-M (Tg(Thy1-eGFP)MJrs/J, Stock No: 007788) was kindly provided by Dr. Yuqiang Ding from Fudan University. ChAT-eGFP (B6.Cg-Tg(RP23-268L19-EGFP)2Mik/J, Stock No: 007902) was kindly provided by Dr. Liang Wang from Zhejiang University.

### Virus injection

Mouse brain injection: The virus injections were conducted following a protocol reported previously (112). A stereotaxic apparatus (RWD Technology Corp., Ltd..) was used to fix the mouse during the injection. A Nanojet III injector (Drummond Scientific, USA) attached to the injection glass micropipette through a plastic tubing was used to inject the virus solution. Virus solutions of rAAV2-cFos-FLPo and rAAV2-CAG-fDIO-eYFP with 1:1 to 1:10 ratios were used for WT mouse injections based on the desired labeling density. Virus solutions of rAAV2-cFos-FLPo and rAAV2-CAG-fDIO-tdTomato with 1:1 to 1:10 ratios were used for ChAT-eGFP mouse injections. The stereotaxic coordinates used to label the spinal tracts originating from different brain areas are as follows. FL: AP +1.2 mm, ML +1.5 mm, DV −0.7 mm; HL: AP −1.2 mm, ML +1.0 mm, DV −0.7 mm; RN: AP −4.0mm, ML +0.75mm, DV −4.0mm; Gi: AP −6.0mm, ML +0.8, DV −5.5. Typically, 100 nl virus solution containing 0.1% Fast Green was injected into the target region of the mouse brain at a rate of 30 nl/min. The spinal cord was dissected ∼6 weeks after the injection.

#### DRG injection

Virus solution of rAAV9-hSyn-Cre and rAAV9-CAG-DIO-tdTomato with 1:1 ratio was used for WT mouse DRG injections, and virus solution of rAAV9-PV-Cre and rAAV9-CAG-DIO-tdTomato with 1:1 ratio was used for ChAT-eGFP mouse DRG injections. Typically, 1 µl virus solution containing 0.1% Fast Green was injected into the target DRG at a rate of 30 nl/min following a protocol reported previously. The spinal cord was dissected ∼6 weeks after the injection.

#### Muscle injection

Virus solution of rAAV9-hSyn-Cre and rAAV9-CAG-DIO-tdTomato with 1:1 ratio was used to label sensory neurons, and virus solution of rAAV2-retro-hSyn-tdTomato was used to label motor neurons. Typically, 1 µl virus solution containing 0.1% Fast Green was injected into the gastrocnemius muscle of the mouse right hindlimb at a rate of 30 nl/min following a protocol reported previously. The spinal cord was dissected ∼6 weeks after the injection.

### Preparation of cleared and expanded mouse spinal cord samples

Spinal cord dissection and fixation: The anesthetized mouse was perfused with heparinized 0.1 M PBS (10 units/ml of Heparin) for 5-10 mins at room temperature until the blood was washed out. Followed by perfusion with pH 7.4 4% paraformaldehyde (PFA) for another 5-10 mins. The mouse spinal cord was dissected following a previously reported protocol (113). The dissected spinal cord was loosely wrapped with a 200 µm Nylon mesh and bound to a perforated Teflon plate with surgical sutures so that the spinal cord can maintain the straight shape through the sample preparation process. The bound spinal cord was placed in 4% PFA overnight at 4°C with gentle shaking.

Preparation of cleared spinal cords (∼5-6 days): The fixed spinal cord was cleared with a modified CUBIC method. First, the fixed spinal cord was immersed in the delipidation solution (15 wt% urea, 10 wt% N-butyldiethanolamine, 10 wt% Triton X-100 and 65 wt% dH_2_O) at 37 °C for three days with gentle shaking. Different hydrophilic delipidation solutions can also be used according to the tissue prosperities, clearing efficiency, cleared tissue transparency and the EF preservation efficiency. The delipidation solution was replaced every 24 hours. Next, the delipidated spinal cord was RI matched by immersing the spinal cord in the RI matching solution of RI ∼1.49 (25 wt% urea, 22.5 wt% sucrose, 22.5 wt% antipyrine, 10 wt% triethanolamine) at 25°C for two days with gentle shaking. The RI matching solution was replaced every 24 hours. Finally, the RI matched spinal cord was embedded in 2% agarose gel made with the RI matching solution following the procedure shown in Fig. 1. The cleared and embedded spinal cord was mounted on the sample holder by attaching the sample to the magnets of the sample holder and imaged with the imaging system using Silicone oil as the imaging buffer.

Preparation of expanded spinal cord samples (∼4 days): The fixed spinal cord was delipidated following the procedure described above. The delipidated spinal cord was washed in 0.01 M PBS for 2-3 hours at room temperature with gentle shaking. The washing buffer was replaced every hour. Next, the washed spinal cord was immerged in the monomer solution (30% Acrylamide (AA), 0.075% N,N-Dimethylacrylamide (BA), 10% Sodium acrylate (SA), 0.5% 2,2’-Azobis[2-(2-imidazolin-2-yl)propane] dihydrochloride (VA-044) in 0.01 M PBS) at 4 °C for 1-2 days with gently shaking. The percent composition of each compound can be adjusted according to the desired expansion ratio as shown in Fig. S1. After the monomer incubation, the incubated spinal cord was polymerized and embedded in the same polymer following the layer-by-layer polymerization procedure shown in Fig. 1. Finally, the polymerized and embedded spinal cord was placed in DI water at 4 °C for ∼2 days for expansion. The DI water was replaced every 12 hours. The expanded and embedded spinal cord or spinal cord sections were glued on a piece of iron sheet and mounted on the sample holder and imaged using DI water as the imaging buffer.

### Microscope

Our imaging system consists of four TLS microscopes of two configurations (Fig. S2). The first configuration was described in our previous publications (65–66). It is capable of imaging centimeter-scale cleared tissues with micron-scale to submicron-scale spatial resolution by using real-time optimized tiling light sheets and detective objectives with various numerical apertures (Olympus XLSLPLN10XSVMP, Nikon Plan Apo 10x Glyc, or Nikon CFI90 20XC Glyc). The resolving ability can be improved to ∼70 nm in combined with tissue expansion. In addition, the microscope is compatible with most tissue clearing and expansion methods and can be aligned semiautomatically through the phase modulation of the illumination light. The microscope conducts multicolor imaging sequentially for up to 5 colors with the excitation wavelengths of 405 nm, 488 nm, 515 nm, 561 nm, and 638 nm.

The other configuration is modified from the first one to conduct simultaneous dual-color imaging with the excitation wavelengths at 488 nm and 561 nm. The 488 nm and 561 nm laser beams are expanded to ∼8 mm (L1=30 mm, L2=250 mm) beam diameter and sent to two identical binary SLM assemblies for phase modulation separately. Each binary SLM assembly consists of a polarizing beam splitter cube and a half-wave plate and a 1280×1024 binary SLM. Each modulated laser beam is focused on an optical slit to block the undesired diffraction orders and combined using a dichroic beam splitter. Both SLMs are conjugated to a galvanometer mirror through relay lenses (L3=300 mm, L4=175 mm). The Galvo mirror directs both illumination beams to one of the two symmetrical illumination paths by offsetting the initial angle of the Galvo mirror and creates two excitation light sheets by scanning the laser beams. The modulated laser beams are further conjugated to the rear pupils of two excitation objectives (Mitutoyo MY5X-802 or MY10X-804) through two pairs of relay lenses (L5=L7=150 mm) separately to illuminate the sample from two opposite directions. The emitted fluorescence is collected with the detection objective (Olympus XLSLPLN10XSVMP), divided with a long-pass dichroic beam splitter, and imaged onto two detection cameras with two tube lenses of identical focal lengths (L=400 mm) after passing two single-band bandpass emission filters. The calibration and operation of the two types of microscopes are nearly the same and described in detail in our previous publications (Chen et al., 2020; Feng et al., 2021).

### Image analysis

The image processing, registration and merging workflow was described in detail in our previous publication (65). The entire workflow was conducted in parallel instead of sequentially in this research. Due to the large data size (typically ∼5-20 TB per sample at the original resolution depends on the imaging volume and number of colors), the 3D image of each subvolume was down sampled for ∼10-100 times after preprocessing, and 3D image of the whole sample was obtained by registering and merging the down sampled 3D images of all subvolumes. Cell counting and segmentation were conducted using Amira semiautomatically. Briefly, a seed mask was created after the denoising, filtering and intensity thresholding of the acquired sample image. The obtained seed mask was inspected and corrected manually. Next, cell segmentation was performed by applying the watershed algorithm to the filtered image with the seed mask. The segmentation result was inspected and corrected again according to the original image for the result. Axon tracing was performed using the Amira filament tracing package manually. For the tracing of axon collateral branches, we traced the branch shaft in Fig. 5, the branch from the node of Ranvier to the furthest branch terminal in Fig. 7 and 7S and the entire branch arbor in Fig. 6S based on the purpose of analysis.

## Acknowledgments

We thank Tian Xu, Ming Yin, Fengquan Zhou, Yongdeng Zhang, Kiryl D. Piatkevich and Bing Zhang for helpful discussions. We thank the advanced biomedical technology core facility, the laboratory animal resource center and the high performance computing center for facility support and technical assistance. L.G. acknowledges support from the Westlake University, the Westlake Education Foundation, the National Natural Science Foundation of China (32150015) and the Zhejiang Province Natural Science Foundation (LR20C070002).

## Author Contributions

L.G. conceived the project and supervised the research; R.F., Y.C., J.L. and L.G. developed the sample preparation methods; D.W. and L.G. constructed the imaging system; R.F., J.X., H.H., Y.C. and L.G. performed the experiments; J.L., R.F., H.H., Y.C., J.X. and L.G. analyzed the data; L.G. wrote the paper with the input of all authors.

## Declaration of Interests

The authors declare no competing interests. Multiple patents regarding the presented imaging system and sample preparation method were filed by Westlake University on behalf of the authors.

## Materials and correspondence

The complete imaging system design, control program, and data analysis program can be obtained for non-commercial use through the technique transfer office of the Westlake University by contacting the corresponding author. The study didn’t generate new reagents or require restricted reagents. Further information and requests for resources and reagents should be directed to and will be fulfilled by the Lead Contact, Liang Gao.

## Notes

https://www.youtube.com/channel/UCSkzm2ENrlMz2OVoFp3EG1Q/videos

