## Supplementary file for "Decoding the mouse spinal cord locomotor neural network using tissue clearing, tissue expansion and tiling light sheet microscopy techniques"

**Supplementary Videos**

Video S1. 3D rendering of a cleared adult Thy1-eGFP mouse spinal cord. Related to Figure 2A.

Video S2. 3D rendering of an expanded adult Thy1-eGFP mouse cervical spinal cord section. Related to Figure 2L.

Video S3. Part 1: 3D rendering of a cleared adult ChAT-eGFP mouse cervical spinal cord section. Related to Figure 3A and 3H. Part 2: 3D rendering of an expanded adult ChAT-eGFP mouse cervical spinal cord ventral horn section. Related to Figure 4L.

Video S4. 3D rendering of the cleared spinal cords of four ChAT-eGFP mice at ages of P1, P7, P28 and P56. Related to Figure 3.

Video S5. 3D rendering of a cleared adult WT mouse spinal cord with sparsely labeled spinal motor neurons, and 3D renderings of 30 segmented spinal cord motor neurons. Related to Figure 4.

Video S6. 3D rendering of an expanded adult ChAT-eGFP mouse lumbar spinal cord section. Related to Figure 4.

Video S7. 3D rendering of a cleared adult WT mouse spinal cord with sparsely labeled sensory neurons. Related to Figure 5.

Video S8. 3D rendering of an expanded lumbar spinal cord section of an adult ChAT-eGFP mouse with sparsely labeled L1 DRG proprioceptive sensory neurons. Related to Figure 5.

Video S9. 3D rendering of a cleared adult WT mouse spinal cord with sparsely labeled FL-spinal CST axons. Related to Figure 6.

Video S10. 3D rendering of a cleared adult WT mouse spinal cord with sparsely labeled HL-spinal CST axons. Related to Figure 6.

Video S11. 3D rendering of an expanded cervical spinal cord section of an adult ChAT-eGFP mouse with sparsely labeled FL-spinal CST axons. Related to Figure 6.

Video S12. 3D renderings of expanded cervical and lumbar spinal cord sections of an adult ChAT-eGFP mouse with sparsely labeled HL-spinal CST axons. Related to Figure 6.

Video S13. 3D rendering of a cleared adult WT mouse spinal cord with sparsely labeled RST axons. Related to Figure 7.

Video S14. 3D rendering of a cleared adult WT mouse spinal cord with sparsely labeled ReST axons. Related to Figure 7 and S7.

Video S15. 3D rendering of an expanded cervical spinal cord section of an adult ChAT-eGFP mouse with sparsely labeled ReST axons. Related to Figure 7.

4K Video YouTube link:

https://www.youtube.com/playlist?list=PLLLHAoLrVoj7G_jB-sXtoJ_QWdDOMumiq

High quality video download link:

https://westlakeu-my.sharepoint.com/:f:/g/personal/gaoliang_westlake_edu_cn/Eu7eON0pbbxMtbk8o0mrrhIBGTM8QjgLS8hDhu83i5lf7g?e=jTzzMs

**Supplementary Figures**

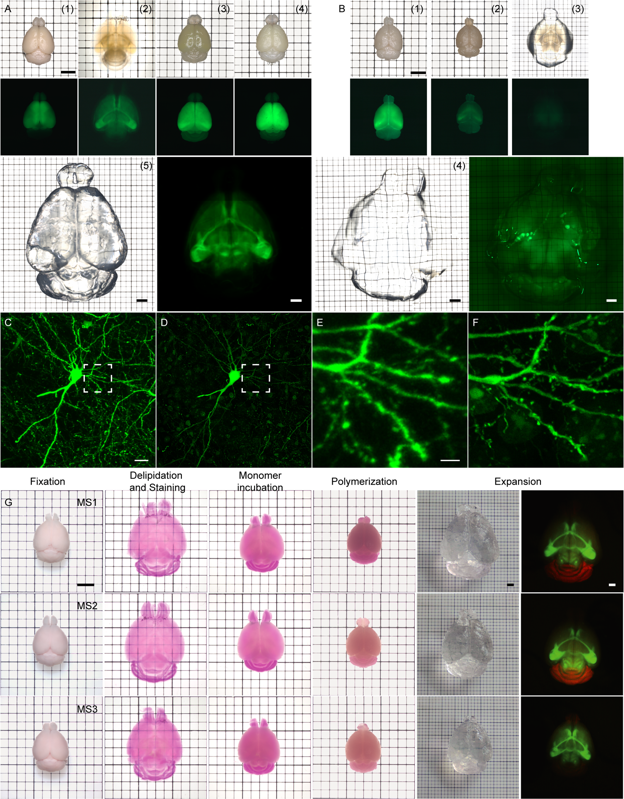

Figure S1. Tissue expansion using CMAP. Related to Figure 1. (A) An adult Thy1-eGFP mouse brain expanded using CMAP. The procedure includes: (1) fixation, (2) delipidation, (3) monomer incubation, (4) polymerization, and (5) expansion. The endogenous fluorophores and tissue structure are well preserved after expansion. (B) An adult Thy1-eGFP mouse brain expanded using MAP. The procedure includes: (1) fixation and monomer perfusion, (2) hybridization and polymerization (45 °C), (3) delipidation and dissociation (70-90 °C), and (4) expansion. The endogenous fluorophores and tissue structure are poorly preserved after expansion. (C, D) The same neuron, within a 100 μm thick Thy1-eGFP mouse brain slice, imaged after delipidation (C) and expansion (D) using CMAP, and the neuronal structure is well preserved despite the elimination of the denaturation and dissociation step. (E, F) Zoom-in views of the selected areas in (C) and (D). (G) Different expansion ratios obtained by changing the percent composition of the monomer solution compounds. MS1: 30% AA 0.1% BA 15% SA 0.5% VA-044; MS2: 30% AA, 0.1% BA, 10% SA, 0.5% VA-044; MS3: 30% AA, 0.1% BA, 5% SA, 0.5% VA-044. Scale bars, 5 mm (A, B and C), 20 μm (D), 5 μm (F).

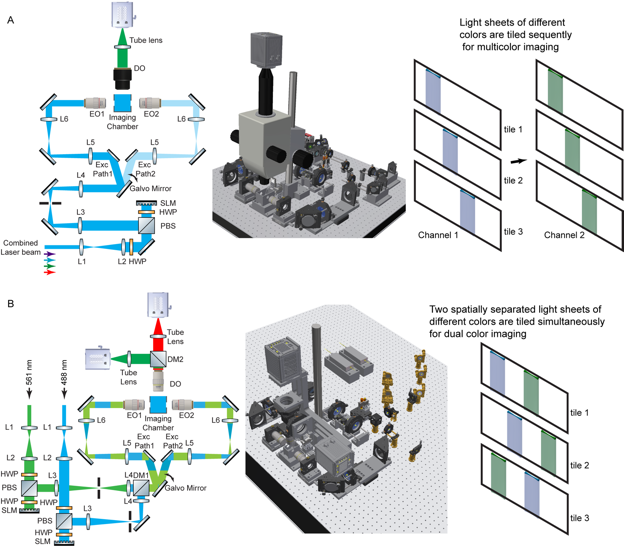

Figure S2. The optomechanical configuration and working principle of the two types of tiling light sheet microscopes implemented for tissue imaging. Related to Figure 1. (A) In the first configuration, tiling light sheets of different excitation wavelengths illuminate the sample sequentially for multicolor imaging for up to five colors. (B) In the second configuration, two independently controlled tiling light sheets with excitation wavelengths of 488 nm and 561 nm illuminate the sample at the same time for simultaneous dual-color imaging.

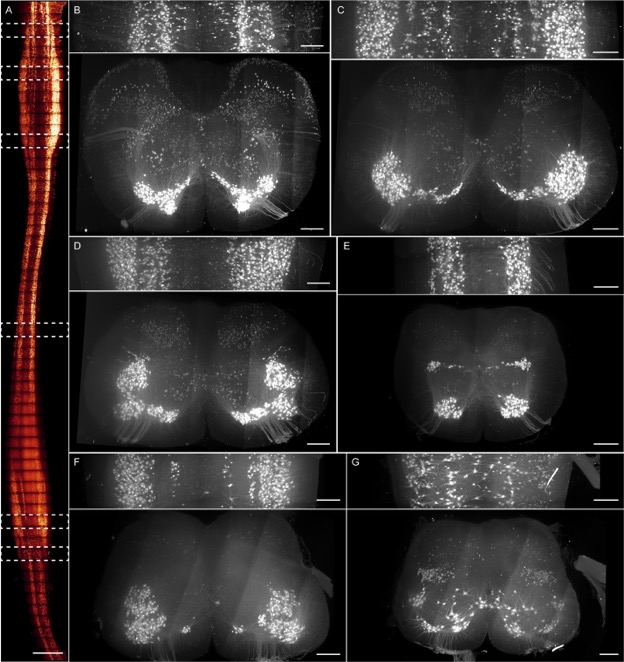

Figure S3. The distribution and dendritic morphology of the SpMNs in various spinal cord sections. Related to Figure 3. (A) Lateral MIPs of a cleared adult ChAT-eGFP mouse spinal cord. (B-G) Lateral and axial MIPs of the selected spinal cord sections in (A), showing the territory of various motor columns. Scale bars, 1 mm (A), 200 μm (B-G).

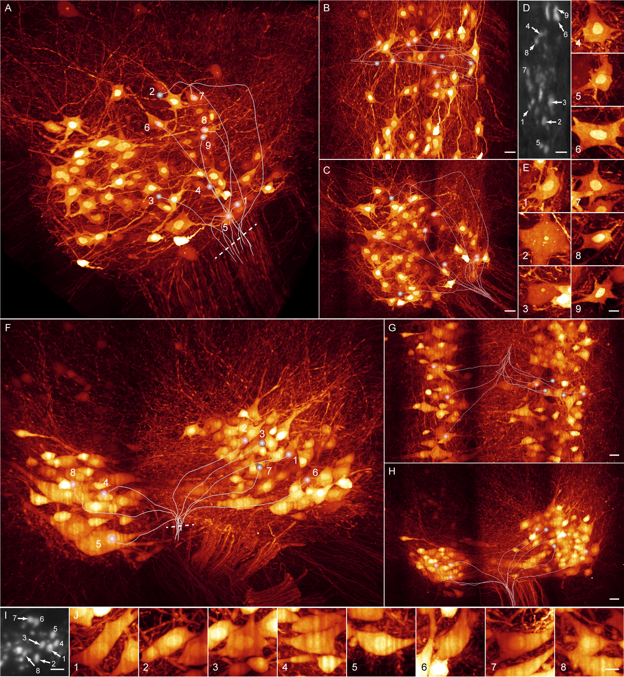

Figure S4. The structure of the mouse spinal cord ventral rootlet. Related to Figure 4. (A-C) 3D rendering and MIPs of an expanded adult ChAT-eGFP mouse thoracic spinal cord ventral horn section that shows the distribution of the SpMNs with axons in the same rootlet. (D) The cross section of the rootlet at the indicated position in (A). (E) Zoom-in views of the traced motor neurons in (A). (F-H) 3D rendering and MIPs of an expanded adult ChAT-eGFP mouse lumbar spinal cord ventral horn section. (I) The cross section of the rootlet at the indicated position in (F). (J) Zoom-in views of the traced SpMNs in (F). Scale bars, 20 μm (B, C, G and H), 10 μm (E and J), 5μm (D and I).

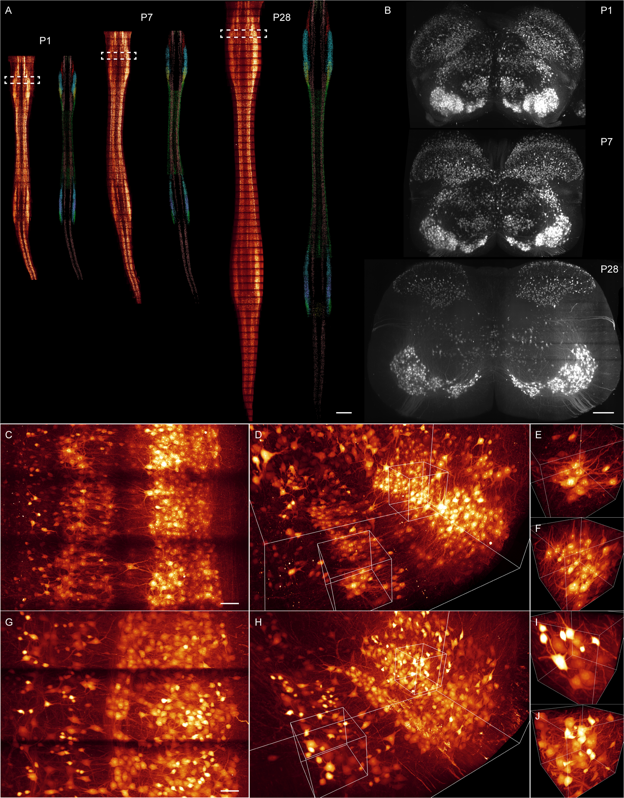

Figure S5. The organization and morphology of the SpMNs in mice of different ages. Related to Figure 3 and 4. (A) Lateral MIPs of the cleared spinal cords of three ChAT-eGFP mice at ages of P1, P7 and P28 and the segregation of the corresponding SpMNs. (B) Axial MIPs of the selected spinal cord sections in (A) showing the different spinal cord motor neuron organizations with the development of mice. (C, D) Lateral MIP and 3D rendering of an expanded P0 ChAT-eGFP mouse cervical spinal cord ventral horn section. (E, F) Zoom-in views of the selected volumes in (D). (G, H) Lateral MIP and 3D rendering of an expanded P56 ChAT-eGFP mouse cervical spinal cord ventral horn section with the location comparable to the section in (C). (I, J) Zoom-in views of the selected volumes in (H). Scale bars, 1 mm (A), 200 μm (B), 50 μm (C and G).

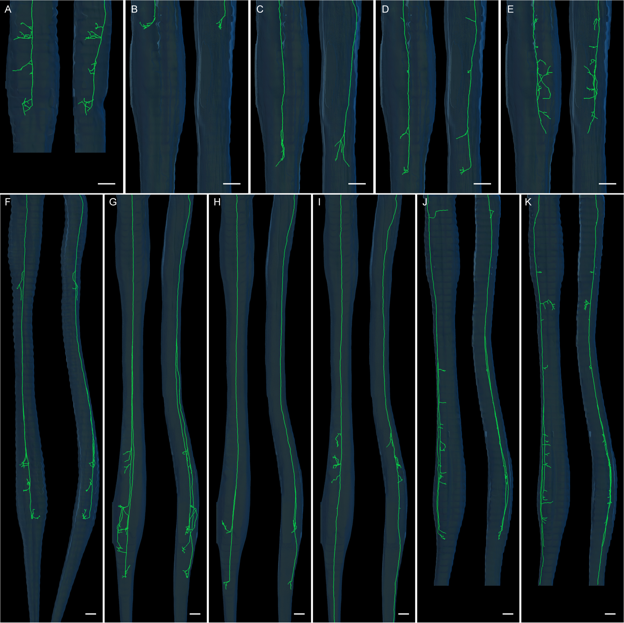

Figure S6. The axonal projection of individual CST axons. Related to Figure 6. (A-E) Lateral and axial MIPs of five FL-spinal CST axons traced form two cleared adult WT mouse spinal cords. Note that the axon (E) project back to the contralateral side (the injection side) of the spinal cord. (F-K) Lateral and axial MIPs of six HL-spinal CSST axons traced form three cleared adult WT mouse spinal cords. Note that a small number of CST axons, such as (J) and (K), descend along the lateral funiculus. Scale bars, 1 mm.

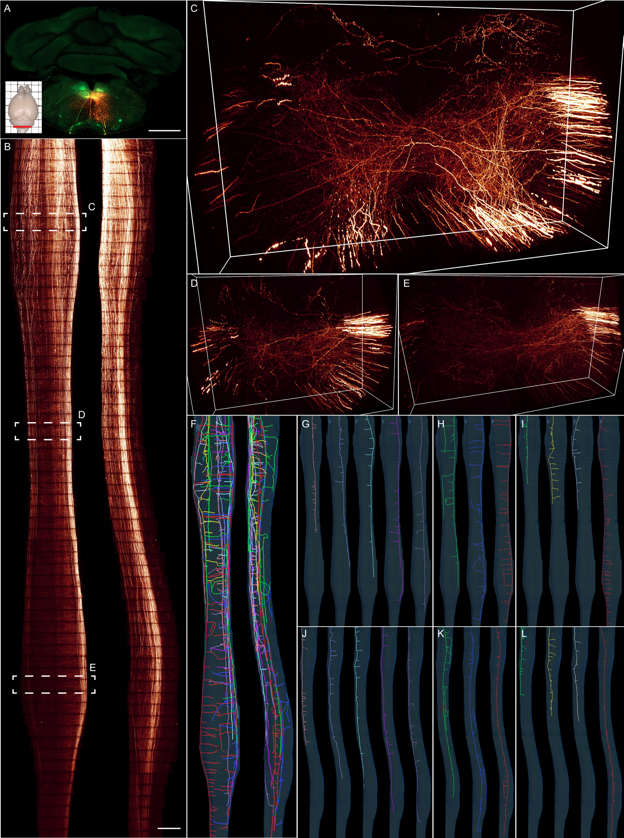

Figure S7. The ReST projection in mouse spinal cord. Related to Figure 7. (A) Sparse labeling of the ReST via Gi virus injection. (B) Lateral and axial MIPs of a cleared adult WT mouse spinal cord with sparsely labeled ReST axons. (C-E) 3D renderings of the selected cervical, thoracic and lumbar sections in (B). (F) The axonal projection of individual ReST axons showing the complicated projection patterns of the ReST. (G-L) Lateral and axial MIPs of the traced individual axons in the ipsilateral ventral (G, H), ipsilateral lateral (I, J), and contralateral ventral funiculus (K, L) respectively. Scale bars, 1 mm.
